# Histone malonylation is regulated by SIRT5 and KAT2A

**DOI:** 10.1101/2022.06.07.495150

**Authors:** Ran Zhang, Joanna Bons, Olga Bielska, Chris Carrico, Jacob Rose, Indra Heckenbach, Morten Scheibye-Knudsen, Birgit Schilling, Eric Verdin

## Abstract

The posttranslational modification lysine malonylation is found in many proteins, including histones. However, it remains unclear whether histone malonylation is regulated or functionally relevant. Here, we report that availability of malonyl-co-enzyme A (malonyl-CoA), an endogenous provider of malonyl groups, affects lysine malonylation, and that the deacylase SIRT5 selectively reduces malonylation of histones. To determine if histone malonylation is enzymatically catalyzed, we knocked down each of the 22 lysine acetyltransferases (KATs) to test their malonyltransferase potential. KAT2A knockdown in particular reduced histone malonylation levels. By mass spectrometry, H2B_K5 was highly malonylated and significantly regulated by SIRT5 in mouse brain and liver. Acetyl-CoA carboxylase (ACC), the malonyl-CoA producing enzyme, was partly localized in the nucleolus, and histone malonylation increased nucleolar area and ribosomal RNA expression. Levels of global lysine malonylation and ACC expression were higher in older mouse brains than younger mice. These experiments highlight the role of histone malonylation in ribosomal gene expression.

## Introduction

Lysine malonylation is a type of protein posttranslational modification (Du et al., 2011; Peng et al., 2011). By adding a negatively charged malonyl group to the positively charged ε-amino group, malonylation increases the space occupied by the modified lysine residue, and alters the electrostatic force around the modified lysine residue. Similar to lysine acetylation, malonylation is reversible, and subject to the removal by SIRT5, a member of class III lysine deacetylases (also known as sirtuins). However, unlike other sirtuin family members, SIRT5 has very weak deacetylase activity. Instead, SIRT5 specifically removes acidic short-chain acyl groups, including malonyl-, succinyl-, and glutaryl-groups, from modified protein substrates (Du et al., 2011; Peng et al., 2011; Tan et al., 2014). Although over a thousand malonylated proteins have been identified to date (Colak et al., 2015; Nishida et al., 2015), their biological significance remains mostly unclear.

Like many other acyl-CoA species, malonyl-CoA, an intermediary metabolite in the de-novo fatty acid synthesis pathway (Saggerson, 2008), is an endogenous provider of malonyl group for lysine malonylation. Acyl-CoA modifications can occur via enzymatic or non-enzymatic pathways. Due to the reactive nature of acyl-CoA thioesters, acylations, including malonylation, are generally believed to be non-enzymatic in mitochondria which provide a basic environment that deprotonates the ε-amino group of lysine and make it more susceptible to nucleophilic attack towards acyl-CoA species (Wagner and Payne, 2013).

In our previous study, we identified 430 malonylated proteins in mouse livers by immuno-affinity enrichment and mass spectrometry (MS) (Nishida et al., 2015). These malonylated proteins were distributed across major subcellular compartments, including mitochondria, cytoplasm, and nucleus (Nishida et al., 2015). Unlike mitochondria, the nucleus has a neutral pH (Casey et al., 2010), and therefore, lysine residues in nuclear proteins are less deprotonated. Meanwhile, malonyl-CoA appears to be less reactive than other acyl-CoA species with similar structures, such as succinyl-CoA and glutaryl-CoA (Wagner et al., 2017). Therefore, within the nucleus, lysine malonylation may need enzymatic catalysis. Yet, no lysine malonyltransferase (KMaT) has so far been reported.

Histones are conserved nuclear proteins with abundant lysine and arginine residues. Using electrostatic attraction, DNA wraps around octamers of the four core histones (e.g., H2A, H2B, H3, and H4) to fit into the nucleus. Histones are subject to chemical modifications, including lysine acetylation, lysine methylation, and arginine methylation, which are some of the best characterized. These modifications play critical roles in regulating DNA replication and gene transcription (Stillman, 2018). With the development of MS techniques in recent years, additional modifications, including lysine malonylation, have been identified in histones (Dai et al., 2020; Du et al., 2011; Peng et al., 2011). However, the functional relevance and regulatory mechanism of histone malonylation are not well understood.

In the present study, we investigated the regulation of histone malonylation. First, we examined the effect of metabolism on lysine malonylation by changing malonyl-CoA levels. Second, we tested the possibility that a known lysine acetyltransferases (KATs) might be able to catalyze histone malonylation (Sabari et al., 2017; Simithy et al., 2017). Third, we tested the effect of histone malonylation on nucleolar expansion and ribosomal RNA expression. Finally, we examined malonylation levels in young and old mouse brains. Through this study, we found that histone malonylation is regulated by malonyl-CoA availability, SIRT5, and KATs. These observations support a role for histone malonylation in aging-associated epigenetic changes.

## Results

### Malonyl-CoA availability affects lysine malonylation

Malonyl-CoA thioester is an endogenous donor of malonyl group for lysine malonylation. Malonyl-CoA is a metabolic intermediate in the de-novo fatty acid synthesis pathway, and is generated by acetyl-CoA carboxylase (ACC). Malonyl-CoA then serves as a substrate for the synthesis of median- and long-chain fatty acids through fatty acid synthase (FAS) (**Figure 1A**). To determine whether lysine malonylation is affected by malonyl-CoA levels, we used compounds that target the metabolism of malonyl-CoA. Using western blotting and an antibody that recognizes malonylated lysine residues (Peng et al., 2011), we examined lysine malonylation in K562 cells at 24 h after treatment with orlistat, a FAS inhibitor). This treatment has been previously reported to increase malonyl-CoA levels (Bruning et al., 2018; Kulkarni et al., 2017). Lysine malonylation was barely detectable without orlistat treatment, and treatment increased the levels of lysine malonylation in a dose-dependent manner (**Figure 1B and 1C**). Next, we tested the effect of ACC inhibition with AICAR. This drug activates AMPK, an inhibitory upstream kinase of ACC through phosphorylation at Ser79 (Davies et al., 1989; Munday et al., 1988; Sullivan et al., 1994). Orlistat treatment significantly increased lysine malonylation, whereas AICAR treatment at 1 mM reduced the orlistat-induced increase in lysine malonylation in K562 cells (**Figure 1D and 1E**). This result indicates that blocking malonyl-CoA biogenesis reduces lysine malonylation.

**Figure 1.**
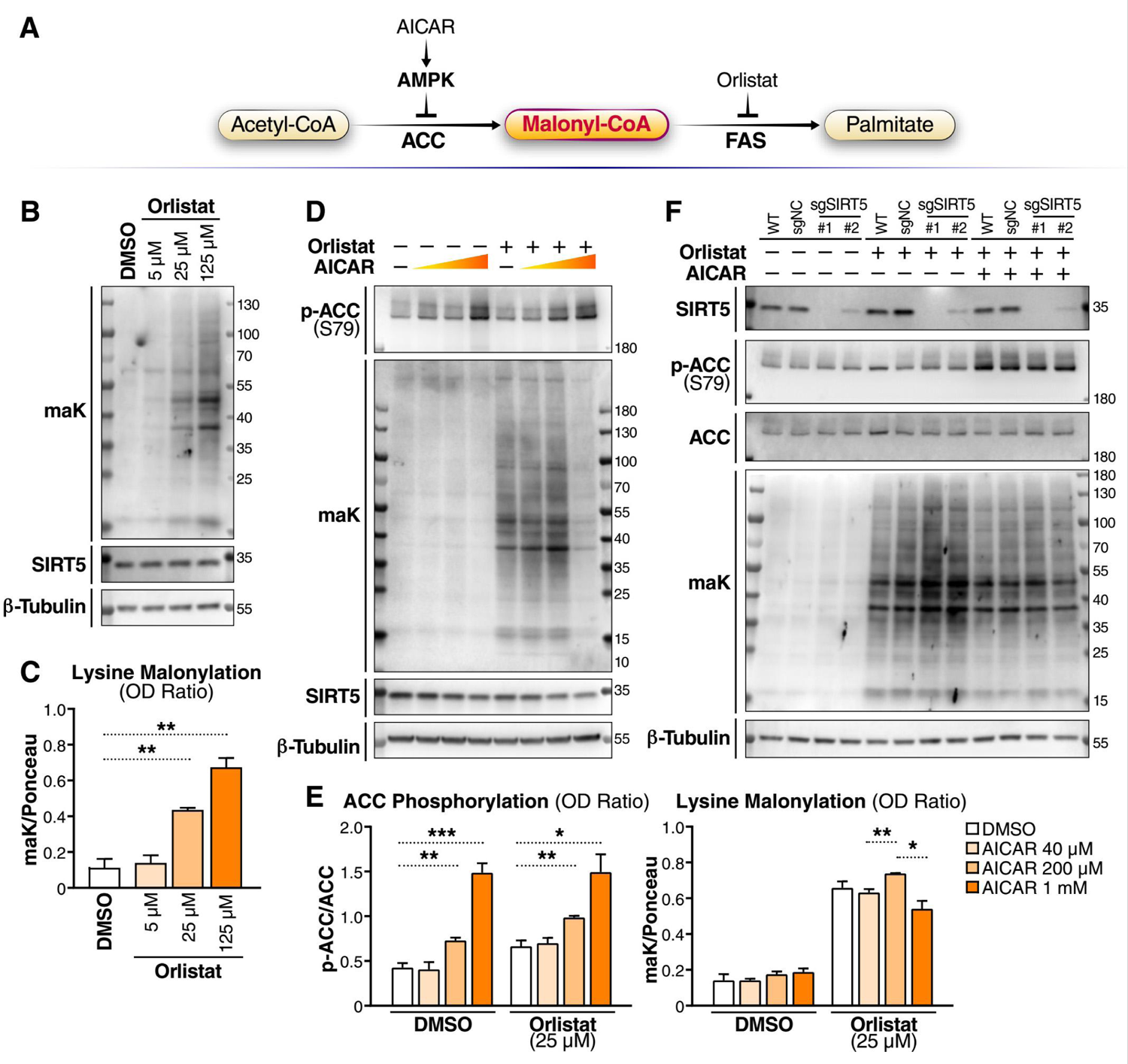
Malonyl-CoA availability affects lysine malonylation. (A) Schematic of the fatty acid synthesis pathway and compounds affecting malonyl-CoA level. AMPK, AMP-activated protein kinase; ACC, acetyl-CoA carboxylase; FAS, fatty acid synthase. (B) Western blot (WB) of lysine malonylation (maK) in K562 cells at 24 h after orlistat treatment at the indicated concentrations. (C) Quantification of WBs by calculating the OD ratio of maK to Ponceau S staining. N = 3. Values are shown as mean ± SEM. **P < 0.01 using unpaired student t test. (D) WB of maK in K562 cells at 24 h after drug treatment. Orlistat: 25 μM; AICAR: 40 μM, 200 μM, or 1 mM. (E) Quantification of WBs by calculating the OD ratio of phosphorylated ACC (at Serine 79) to total ACC, and OD ratio of maK to Ponceau S staining. N = 3. Values are shown as mean ± SEM. *P < 0.05, **P < 0.01, and ***P < 0.001 using unpaired student t test. (F) WB of maK in K562 (wildtype) cells, and K562-dCas9-KRAB cells expressing sgNC, sgSIRT5#1 or sgSIRT5#2 at 20 h after drug treatment. Orlistat: 25 μM; AICAR: 1 mM.

SIRT5 is a prominent deacylase that removes malonyl from lysine residues. To determine the effect of SIRT5, we modified K562 cells by knocking down SIRT5 using a CRISPRi system (Gilbert et al., 2014) with two different guide RNAs (gRNAs) targeting the SIRT5 promoter. Lysine malonylation was barely detected in both control (wildtype (WT) and sgNC (expressing scramble gRNA as negative control)) and both SIRT5 knockdown (KD) K562 cells without orlistat treatment (**Figure 1F**). After orlistat treatment at 25 μM for 24 h, lysine malonylation was induced, and as expected, lysine malonylation levels were higher in SIRT5 KD cells than controls (**Figure 1F**). However, following inhibition of ACC activity with AICAR, lysine malonylation levels in SIRT5 KD cells were no longer higher than in control cells (**Figure 1F**). This result suggests that SIRT5 lowers the hypermalonylation induced by the blockage of malonyl-CoA consumption (FAS inhibition), and the demalonylation function of SIRT5 becomes less effective when malonyl-CoA biogenesis is also reduced in K562 cells. Overall, these results indicate that lysine malonylation is affected by SIRT5 and malonyl-CoA availability.

### SIRT5 and lysine acetyltransferases regulate histone malonylation

Next, we tested whether histone proteins are malonylated. Histones are basic nuclear proteins with abundant lysine residues that are subject to multiple acylations (Huang et al., 2014). We extracted histones from WT and SIRT5 knockout (S5KO) mouse liver, and examined their acylation levels using pan-lysine-acylation antibodies. Western blotting result showed that lysine acetylation was abundant in WT and S5KO mouse histones (**Figure 2A**). In agreement with the known low deacetylase activity of SIRT5 (Du et al., 2011), acetylation levels were not different in WT and S5KO histone samples (**Figure 2A**). Since SIRT5 functions mainly as a lysine demalonylase, desuccinylase, and deglutarylase (Du et al., 2011; Peng et al., 2011; Tan et al., 2014; Zhang et al., 2011), we probed lysine malonylation, succinylation, and glutarylation in the histone samples. Western blotting showed that lysine malonylation was barely detected in WT mouse liver histone samples, but was remarkably higher in S5KO mouse livers (**Figure 2A**). However, lysine succinylation and glutarylation levels were very low in histones, and no significant differences were observed in WT and S5KO mouse histone samples (**Figure 2A**). These results suggest that histones are more susceptible to malonylation than succinylation or glutarylation in the absence of SIRT5.

**Figure 2.**
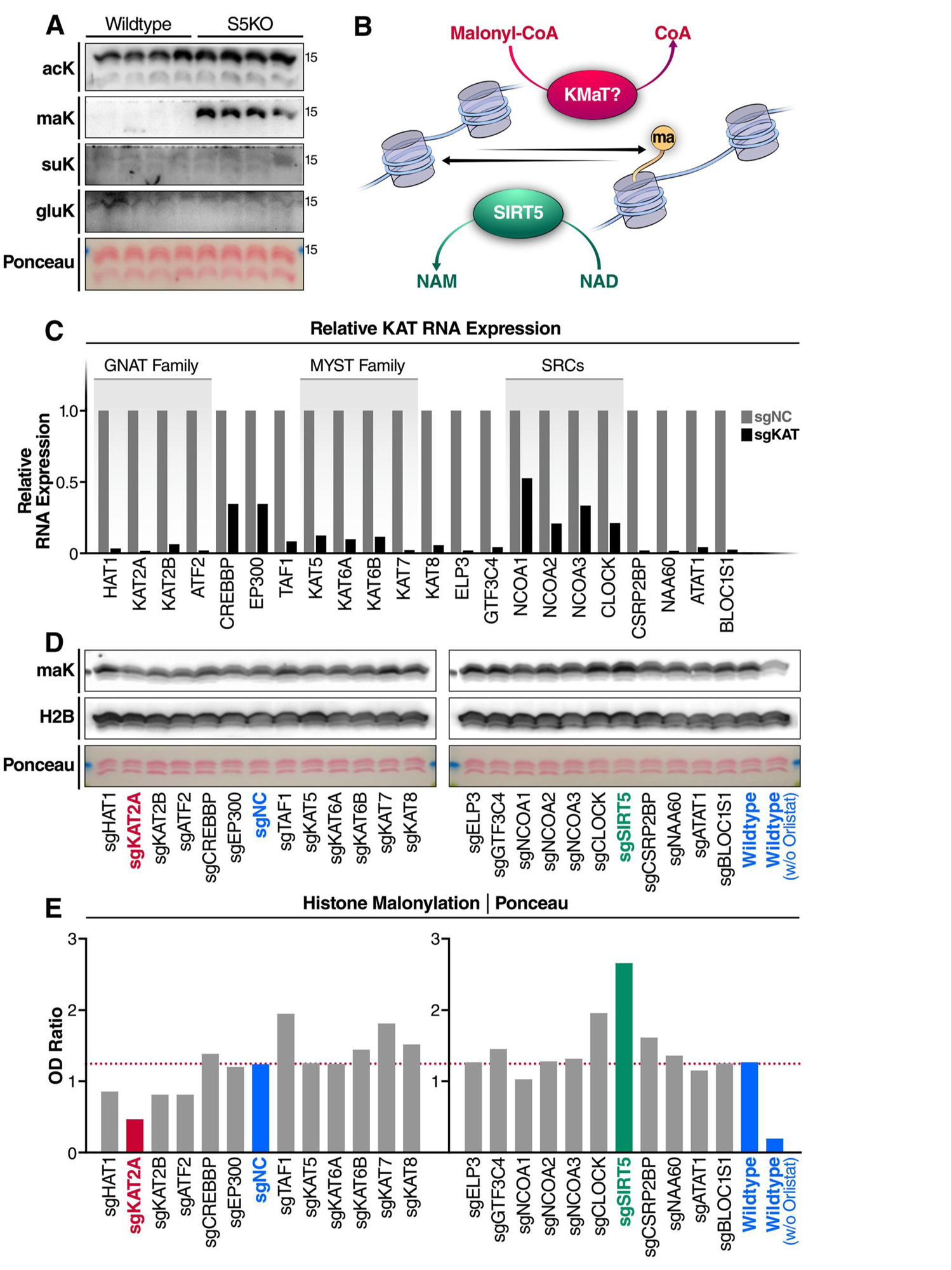
SIRT5 and lysine acetyltransferases regulate histone malonylation. (A) Western blot (WB) of acetylation, malonylation, succinylation, and glutarylation in histone extracts derived from wildtype and SIRT5 knockout (S5KO) mouse livers. (B) Schematic of the regulation of histone malonylation. KMaT, lysine malonyl-transferase. (C) Lysine acetyltransferase (KAT) knockdown K562 cell lines were constructed using a CRISPR-dCsa9-KRAB system via lentiviral infection. Knockdown efficiencies were examined by qPCR. (D) WB examination of malonylation on histones extracted from KAT knockdown K562 cells treated with 25 μM orlistat for 24 h WT cells without lentiviral infection, or with scramble lentiviral infection (sgNC), as control cell lines (shown in blue). (E) Quantification of the optical densities (OD) of malonylation blots normalized to Ponceau S staining.

We next wanted to know if histone malonylation might be regulated by a lysine malonyltransferase (KMaT) (**Figure 2B**). Lysine acetyltransferases (KATs) transfer acetyl groups from acetyl-CoA to lysine residues. In the human proteome, 22 putative KATs have been identified so far. With some exceptions, KATs are mainly catalogued into three families (i.e., GCN5-related N-acetyltransferases (GNAT), the p300/CREB-binding protein (p300/CBP), and the MOZ, Ybf2, Sas2, and Tip60 (MYST) family) (Ali et al., 2018). Importantly, KATs are not exclusive acetyltransferases; some also catalyze non-acetyl lysine acylations (Sabari et al., 2017).

Using a CRISPRi system (Gilbert et al., 2014), we knocked down each of the 22 KATs in K562 cells, and tested their malonyl-transferring potential. For each KAT gene, we used two different gRNAs with the highest predicted targeting potentials, and then chose the one with higher knockdown efficiency in K562-expressing dCas9-KRAB (Krüppel-associated box) cells to evaluate histone malonylation (**Table S1**). RNA expression levels of different KAT genes upon CRISPRi KD were reduced by 47–99%, and 18 of 22 KAT genes had a knockdown efficiency > 80% (**Figure 2C**). These sgKAT-expressing K562 cells were then treated with 25 μM orlistat for 24 h before histone extraction. Western blotting using a pan-malonyllysine antibody detected lysine malonylation in histones in different sgRNA-expressing K562 cells, and, as predicted, malonylation levels were much lower in K562 cells without orlistat treatment (**Figure 2D**, right most lane). WT (not expressing gRNA) and sgNC K562 cells had comparative levels of histone malonylation (**Figure 2D and 2E**). As expected, histone malonylation level in SIRT5 KD cells (shown in green) was higher than in WT or sgNC cells (**Figure 2D and 2E**). Western blotting results also showed that some KAT KD cells (e.g., sgTAF1, sgKAT7, and sgCLOCK) had higher, whereas some KAT KD cells (e.g., sgHAT1, sgKAT2A, sgKAT2B, sgATF2) had lower histone malonylation levels than WT or sgNC cells (**Figure 2D and 2E**). Of particular interest, among all KAT KD K562 cells, the knockdown of KAT2A (shown in red) caused the highest level of reduction (by > 60%) in histone malonylation compared to WT or sgNC cells (**Figure 2D and 2E**), which suggests KAT2A as a robust regulator of histone malonylation.

To confirm the observation that KAT2A KD reduces histone malonylation in the CRISPRi screening experiment, we examined histone malonylation levels in two KAT2A KD K562 cell lines that express two different KAT2A-targeting sgRNAs. As shown in **Figure 3A and 3B**, after orlistat treatment, two different SIRT5 KD K562 cell lines both had 50–100% higher levels of histone malonylation, whereas two different KAT2A KD cell lines had lower levels (decreased by ∼50%). These results confirmed that knockdown of KAT2A reduces orlistat-induced histone malonylation. Then we made a SIRT5 and KAT2A double knockdown K562 cell line and examined its histone malonylation levels. After orlistat treatment, knockdown of SIRT5 alone significantly increased histone malonylation, whereas knockdown of KAT2A significantly reduced histone malonylation in SIRT5 KD cells (**Figure 3C and 3D**), indicating that histone malonylation is reversible and regulated by both SIRT5 and KAT2A. These results confirmed that KAT2A regulates histone malonylation and contributes to orlistat-induced histone malonylation.

**Figure 3.**
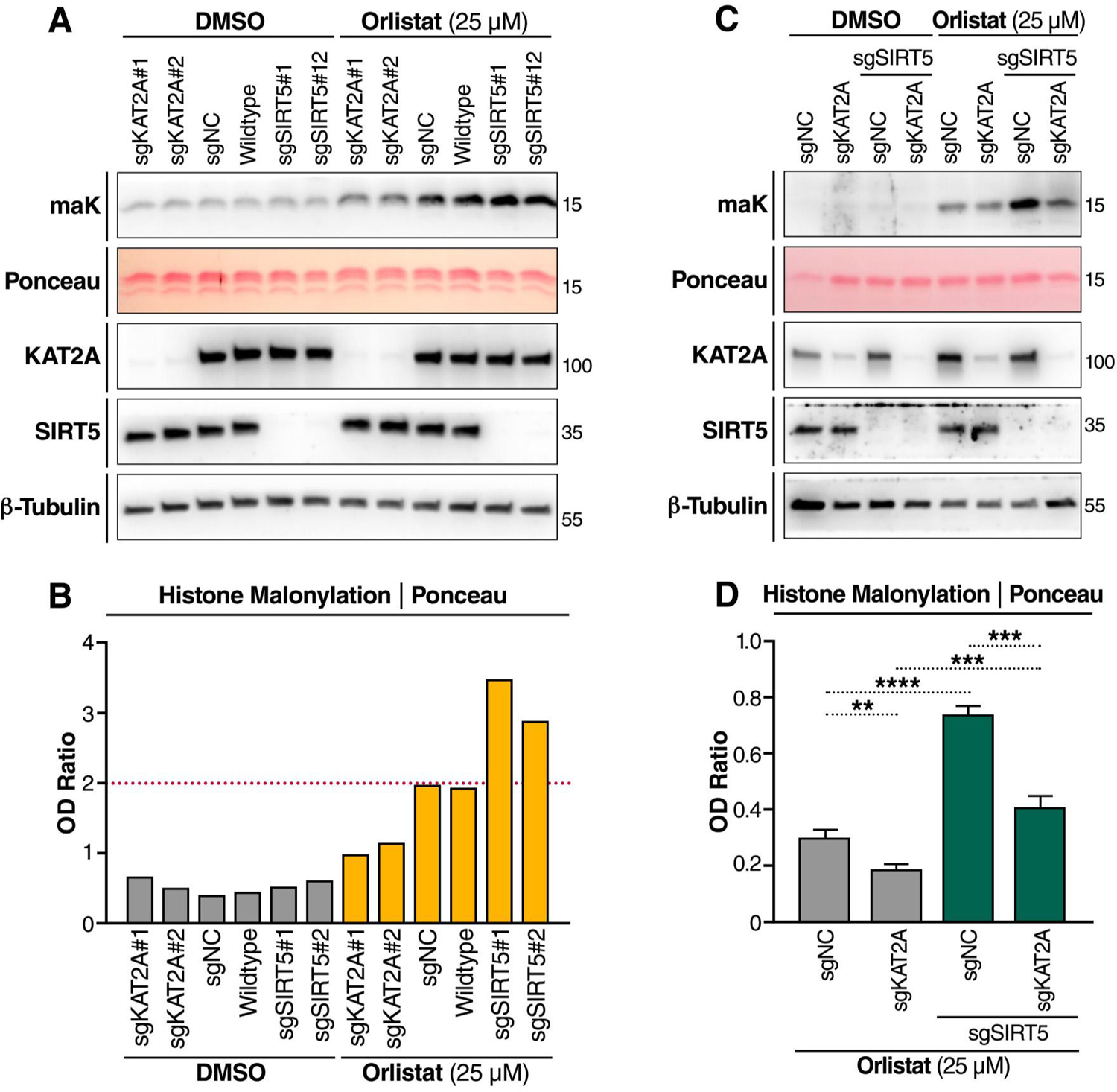
KAT2A regulates histone malonylation. (A) Western blot (WB) of malonylation in histone extracts. sgKAT2A (two different sgRNAs), sgNC, WT, and sgSIRT5 (two sgRNAs) K562 cell lines were treated with 25 μM orlistat or DMSO for 24 h. (B) Optical density (OD) quantification of histone malonylation normalized to Ponceau S staining in (A). (C) WB of malonylation in KAT2A single knockdown and SIRT5;KAT2A double knockdown K562 cells. (D) OD quantification of histone malonylation normalized to Ponceau S staining. n = 4 per cell type. Error bar: mean ± SEM. **P < 0.01, ***P < 0.001, and ****P < 10^-4^ using unpaired student t test.

To identify the lysine sites in histones that are actually malonylated *in vivo*, we examined the malonylation proteome in mouse tissues. We used MS and data-independent acquisitions (Collins et al., 2017; Gillet et al., 2012) to profile the malonylation proteome in mouse brain after immunoaffinity enrichment of malonylated peptides. H2B_K5 was highly malonylated in SIRT5 KO (S5KO) mouse brains, but not detectable in WT mouse brains (Log_2_(S5KO vs WT) = 9.21, with p-value < 0.0001) (**Figure 4A and 4B**), suggesting that H2B_K5 is highly malonylated and regulated by SIRT5. To confirm this observation, we went back to our previous study, in which we probed malonylated proteins in WT and S5KO mouse liver tissue lysates by MS (Nishida et al., 2015). Re-analysis of the data set identified nine malonylated lysine sites in histones in mouse liver (**Figure 4C**). Like mouse brain, liver showed H2B_K5 malonylation levels also higher in S5KO vs. WT mouse livers (**Figure 4C**). These results suggest that H2B_K5 malonylation is regulated by SIRT5.

**Figure 4.**
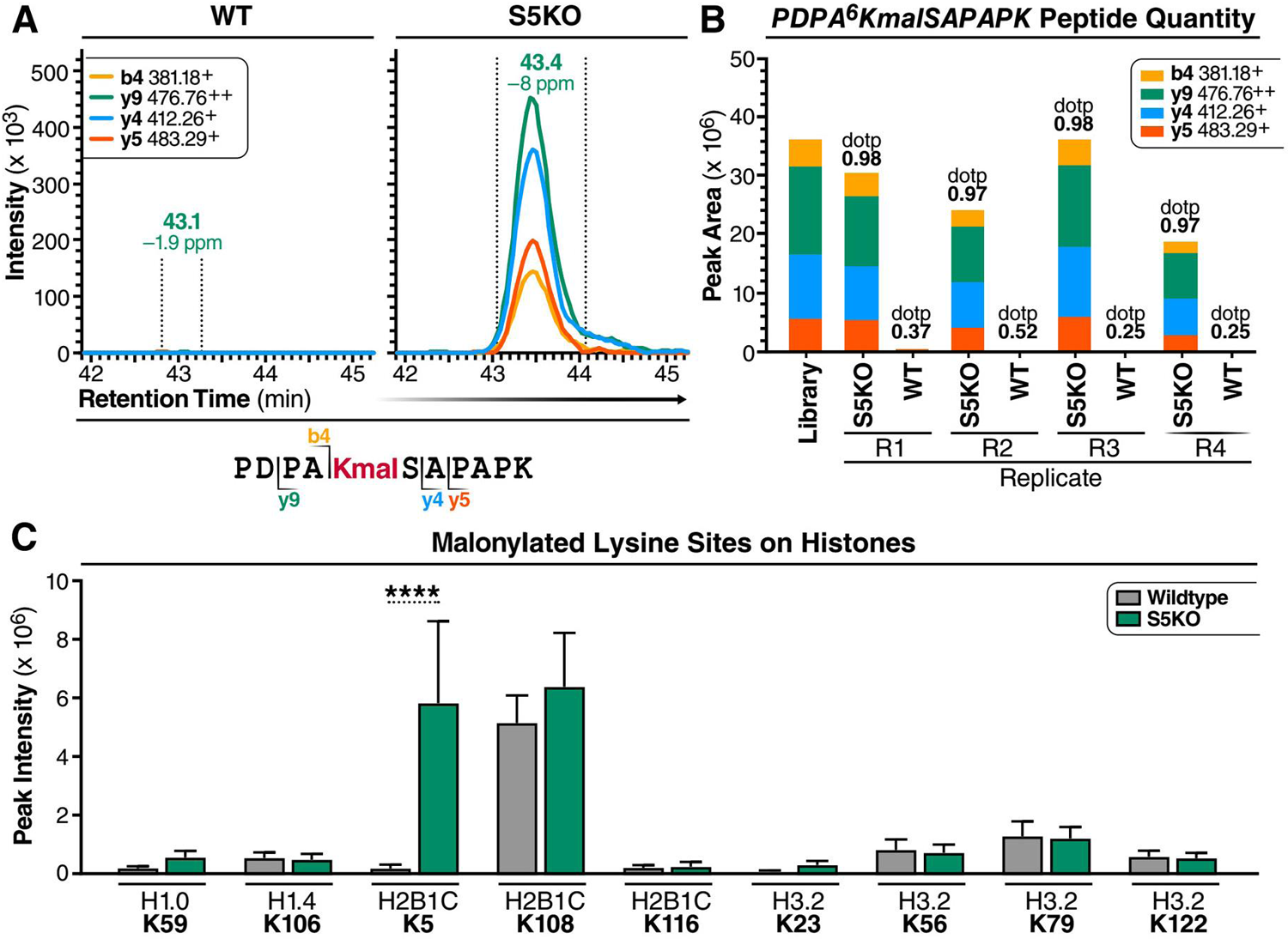
Histone H2B is malonylated at K5 in SIRT5 KO mouse brain and liver. (A) Extracted ion chromatograms of PDPA^6^KmalSAPAPK (m/z 589.81, z=2+) from mouse histone H2B type 2-B (Q64525) obtained by data-independent acquisition, that indicates that malonylation of K5 is increased in S5KO mouse brain tissues compared to wildtype (WT). (B) Quantification using the peak areas of four transitions of PDPA^6^KmalSAPAPK (m/z 589.81, z=2+) in four biological replicates of S5KO and WT mouse brain tissues, respectively. (C) Malonylated lysine sites on histones were previously identified in WT and S5KO mouse livers (Nishida, et al., Mol Cell. 2015). n = 5. Error bar: mean ± SD. ****P < 10^-4^ using Sidak multiple comparisons test following two-way ANOVA.

### Histone malonylation increases ribosomal RNA expression

Next, we tested a possible biological role for histone malonylation. Since lysine malonylation is affected by local availability of malonyl-CoA, the localization of malonyl-CoA-producing enzyme, ACC, should inform the biological function of malonylation. Two isoforms of the ACC protein (i.e., ACC1 and ACC2) are expressed in mammals. ACC1 mainly localizes to the cytoplasm, and ACC2 localizes to both cytoplasm and the outer membrane of mitochondria (Tong and Harwood, 2006). To examine the localization of ACC1, we used western blotting to examine ACC expression in subcellular fractions of HEK293T cells. Although ACC was expressed highest in the cytosolic fraction, a low level of ACC was detected in the nuclear fraction (**Figure 5A**), suggesting the nucleus as an unconventional subcellular compartment for ACC1. Nuclear localization of ACC1 is also reported in the human protein atlas (HPA) database (proteinatlas.org), which shows that ACC1 (also known as ACACA) was identified in nucleoli (**Figure S1**). To confirm this result, we performed confocal imaging after immunofluorescence staining of ACC and fibrillarin (FBL), a nucleolar marker, in HeLa cells. By fluorescence distribution analysis, we observed that ACC was detected in the nuclei, including nucleoli (where FBL was stained), in HeLa cells (**Figure 5B and 5C**). Hence, ACC1 may function as a local malonyl-CoA provider, thereby increasing malonyl-CoA availability and histone malonylation in nucleoli.

**Figure 5.**
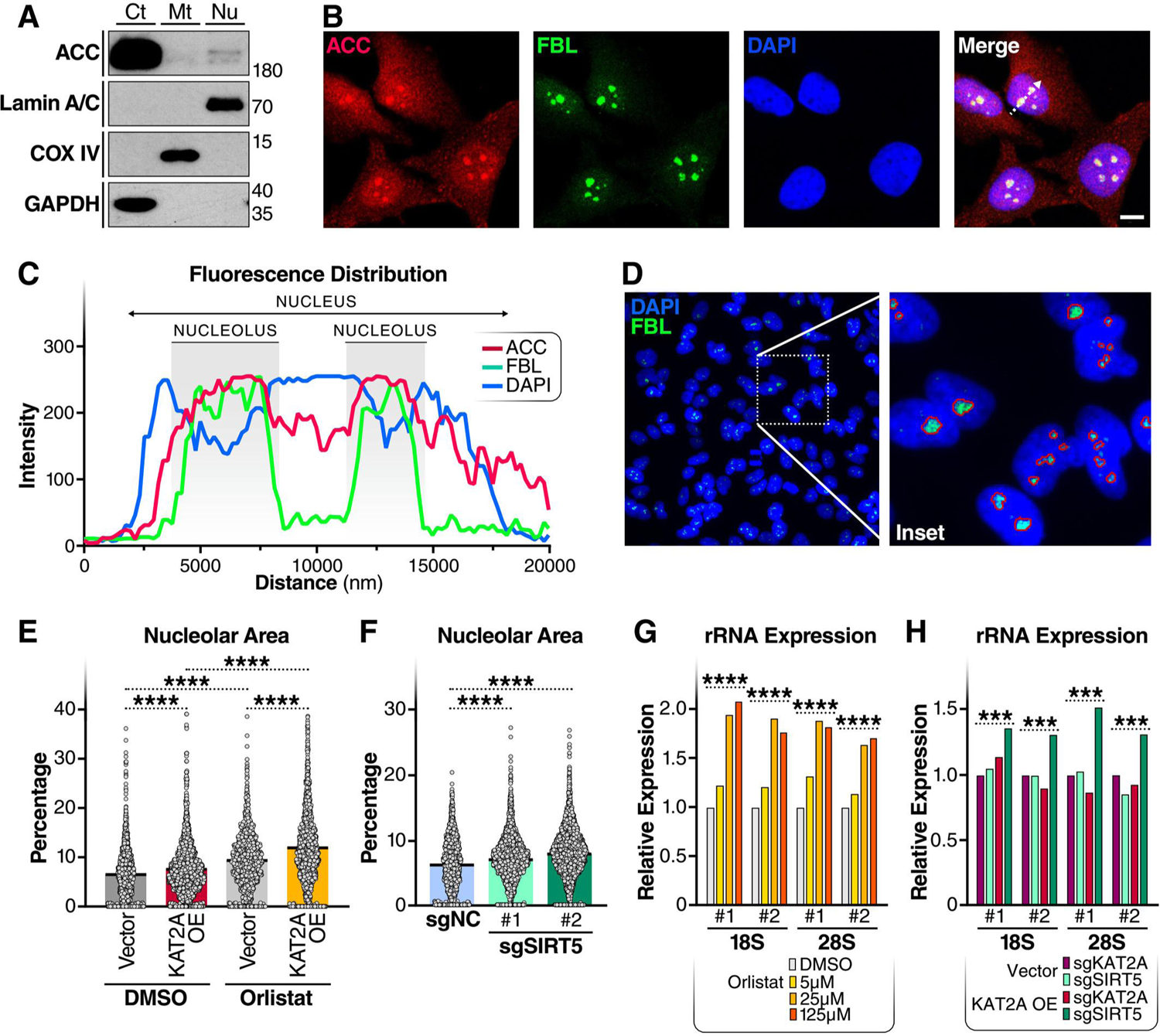
Lysine malonylation increases ribosomal RNA expression. (A) WB with subcellular fractions of HEK293T cells. Ct, cytosol; Mt, mitochondria; Nu, nucleus. (B) Representative image of immunofluorescence staining and confocal imaging of ACC (red), fibrillarin (FBL, green), and DAPI (blue) in HeLa cells. Scale bar: 10 μm. White dotted arrow area in the overlay (Merge) image was sampled for fluorescence distribution analysis in (C). (C) The distribution of fluorescence signals of ACC (red), FBL (green), and DAPI (blue) along the dotted arrow in the overlay image in (B) was plotted to show the relative localizations of ACC, FBL, and DAPI. (D) Representative image of immunofluorescence staining and wide-field imaging of FBL (green) and DAPI (blue) in HeLa cells using a 40x objective. Inset shows nucleolar areas identified (circled with red lines) using a machine learning algorithm. (E) Nucleolar percentage area (ratio of nucleolar area to nuclear area) was quantified in HeLa cells transfected with empty vector (Vec) or KAT2A expressing plasmid (KAT2A OE) 24 h before DMSO or orlistat treatment (25 μM) for another 24 h. N = 1201–2265. ****P < 10^-4^ using unpaired student t test. (F) Nucleolar percentage area was quantified in HEK293T cells expressing MeCP2-KRAB-dCas9 with scramble guild RNA (sgNC) or two different gRNAs targeting SIRT5 promoter (sgSIRT5#1 and #2). N = 1352–3054. ****P < 10^-4^ using unpaired student t test. (G) qPCR examination of ribosomal RNA levels in HeLa cells after DMSO or orlistat treatment for 24 h. rRNA expression levels were normalized to β level. ****P < 10^-4^ using Sidak multiple comparisons test following two-way ANOVA. (H) qPCR examination of ribosomal RNA levels in sgKAT2A, sgSIRT5 K562 cells upon vec or KAT2A overexpression (O/E). rRNA expression levels were normalized to β-actin expression level. ***P < 0.001 using Sidak multiple comparisons test following two-way ANOVA.

The malonyl group is bulkier than the acetyl group, and malonylation changes the positive charge at the lysine side chain to a negative charge might play a role in chromatin compaction in nucleoli which might be reflected in nucleolar size. To test this hypothesis, we measured nucleolar size using immunofluorescence staining and a machine-learning algorithm to quantify nucleolar area (as a percentage of nuclear area) in HeLa cells (**Figure 5D**). Cells treated with 25 μM orlistat for 24 h had larger nucleolar percentage areas than DMSO-treated cells (**Figure 5E**). KAT2A overexpression increased nucleolar percentage area in both DMSO or orlistat treated HeLa cells (**Figure 5E**). Similarly, we quantified nucleolar areas in SIRT5 KD HEK293T cells. The nucleolar percentage areas were significantly higher in SIRT5 KD cells than in control cells (**Figure 5F**). These results support the notion that inducing histone malonylation increases nucleolar area.

Since ribosomal RNAs (rRNAs) are transcribed and produced in the nucleoli, we next examined rRNA expression levels by qPCR in these cells. Orlistat treatment increased the expression levels of both 18S and 28S rRNAs in a dose-dependent manner in HeLa cells (**Figure 5G**). Similarly, we examined rRNA expression levels in K562 cells treated with 25 μM orlistat for 24 h. Overexpressing KAT2A increased the expression levels of both 18S and 28S rRNAs in SIRT5 KD cells, but not in KAT2A KD cells (**Figure 5H**), suggesting that the KD of SIRT5 is required for the induction of rRNA expression upon KAT2A overexpression. These results indicate that lysine malonylation increases rRNA expression.

### Lysine malonylation increases with age in mouse brain

In light of the observations and adverse effect of nucleolar expansion in both premature and physiological aging (Buchwalter and Hetzer, 2017; Tiku et al., 2016), we tested whether lysine malonylation could be involved in the aging process. We extracted protein lysates from young (2 months old) and older (18 months old) mouse brains, and examined lysine malonylation levels by western blotting. Global malonylation levels, including malonylation of proteins around 15 kDa, where histones are located (shown upon long exposure in **Figure 6A**), were higher in older than in young mice (**Figure 6A and 6B**), suggesting that levels of lysine malonylation increase with aging. Expression levels of SIRT5 and KAT2A were not different in young and older mice (**Figure 6A and 6B**), but the levels of ACC, the enzyme that generates malonyl-CoA, were higher in brains of older mice (**Figure 6A and 6B**), suggesting that older mice may have a higher level of fatty acid synthesis in the brain that contributes to the increase in lysine malonylation.

**Figure 6.**
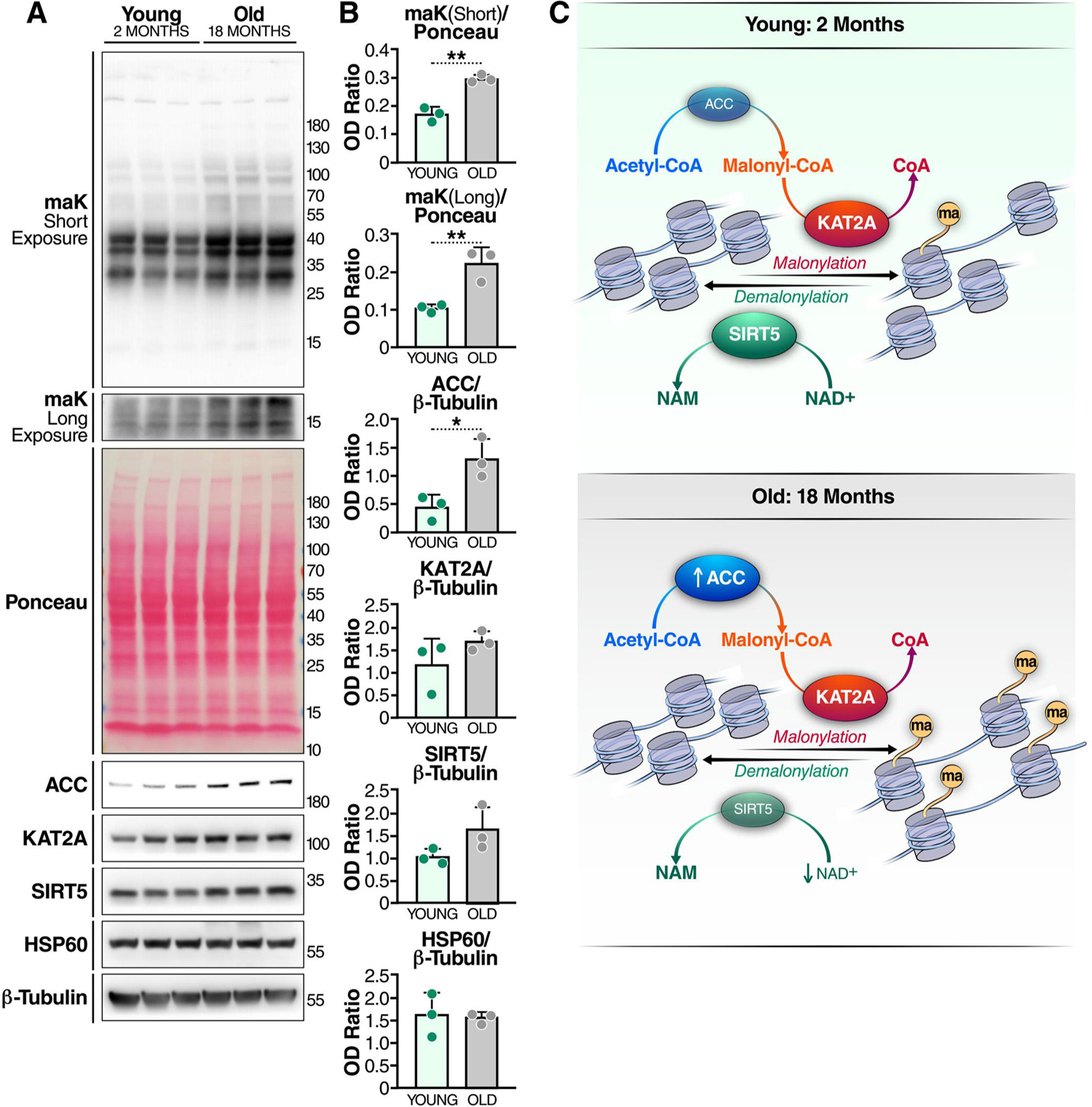
Histone malonylation as a function of age. (A) Western blot of protein samples prepared from young (2 months old) and middle-aged (18 months old) female C57BL/6 mouse brains. (B) Quantification of the ODs of blots. n = 3 per group. Error bar: mean ± SD, *p<0.05, **p<0.01 with unpaired student t test. (C) Schematic of histone malonylation regulated by malonyl-CoA level, KAT2A, and SIRT5 emerges as a mechanistic link connecting metabolism, gene expression, and aging.

As a sirtuin family member, the deacylase activity of SIRT5 requires the availability of co-substrate NAD^+^ (Houtkooper et al., 2010). NAD levels decline with age (Lautrup et al., 2019; Zhu et al., 2015), thereby restraining the activities of sirtuins, including SIRT5, at older age. Although the western blotting showed that the protein level of SIRT5 was not significantly different in young and older mouse brains (**Figure 6A and 6B**), a decreased SIRT5 activity might still be a contributing factor to the increased level of lysine malonylation in old mice. Overall, these results suggest a possible association between histone malonylation and aging (**Figure 6C**).

## Discussion

In the present study, we showed that lysine malonylation is affected by malonyl-CoA availability and found that histone malonylation is regulated by SIRT5 and KAT2A. Importantly, we observed that rRNA expression is subject to the regulation by lysine malonylation, and that global malonylation increases with age in mouse brain. Hence, this study reveals a possible role for histone malonylation in epigenetic regulations.

Malonyl-CoA thioester is a moderately reactive metabolic intermediate in *de-novo* fatty acid synthesis pathway that provides malonyl group to modify free lysine residues. Genetic ablation of malony-CoA consuming enzymes, including MCD and FAS, globally increases lysine malonylation (Bruning et al., 2018; Colak et al., 2015), whereas knocking out malonyl-CoA synthetase, ACSF3, reduces lysine malonylation in mitochondria (Bowman et al., 2017). Thus, lysine malonylation appears to be tightly linked to malonyl-CoA metabolism. We confirmed the importance of malonyl-CoA availability in regulating lysine malonylation through pharmacological manipulation of the malony-CoA metabolic pathway. The basal level of lysine malonylation is barely detectable, even upon SIRT5 KD, in K562 cells, possibly due to constant proliferation and lack of malonylation accumulation in cultured cells. However, as shown here and elsewhere (Kulkarni et al., 2017), lysine malonylation can be highly induced by orlistat (FAS inhibitor) treatment in cultured cells, and knockdown of SIRT5 further increased the level of orlistat-induced global lysine malonylation. On the other hand, orlistat-induced malonylation can be reduced by inhibiting ACC activity by activating AMPK using AICAR at a high dosage (1 mM). These observations suggest lysine malonylation as a function of malonyl-CoA metabolism.

SIRT5 is the main deacylase targeting malonyl, succinyl, and glutaryl groups for removal (Du et al., 2011; Peng et al., 2011; Tan et al., 2014). In the present study, we found that levels of histone malonylation are dramatically increased in SIRT5 KO mouse liver, confirming that SIRT5 is a histone demalonylase. However, no difference in the overall histone succinylation or glutarylation was detected upon SIRT5 KO, possibly due to low levels of succinylation and glutarylation in histones. This result is not entirely surprising since succinyl-CoA is a TCA cycle intermediate, and glutaryl-CoA is an intermediate in amino acid metabolism, and both metabolites are generated in mitochondria rather than in the nucleus. However, malonyl-CoA is a fatty acid building block generated in cytoplasm and, possibly, in nucleus (regarding the nuclear localization of ACC), which allows it to modify histones. By MS, we further determined that histone H2B was highly malonylated at K5 in both brain and liver tissues in SIRT5 KO mice. Although H2B_K108 was also identified to be highly malonylated in mouse liver, it appears that the malonyl group on H2B_K108 was not subject to the removal by SIRT5, possibly due to the lack of accessibility of H2B_K108 for SIRT5. Hence, these results suggest that H2B_K5 malonylation is selectively regulated by SIRT5.

With the discovery of a growing list of lysine acylations, KATs have been reported to possess an expanding repertoire of acyltransferase activities (Sabari et al., 2017). Besides lysine acetylation, EP300 catalyzes propionylation, butyrylation, β-hydroxybutyrylation in histones (Chen et al., 2007; Huang et al., 2021; Kaczmarska et al., 2017; Sabari et al., 2015). Structural studies suggest that EP300 contains a deep aliphatic pocket within its active site that accommodate a wide range of acyl-CoA species (Kaczmarska et al., 2017). However, during the screening of KAT KD K562 cells for histone malonyltransferase, we observed no difference in histone malonylation upon EP300 knockdown. This discrepancy might be due to several reasons: 1) relatively low knockdown efficiency (65%) of EP300 in K562 cells, 2) high selectivity of EP300-regulated histone malonylation on the chromatin, or 3) redundancy of malonyltransferase activity between EP300 and other KATs, such as CBP. By contrast, we observed that KD of KAT2A strongly decreased histone malonylation. This observation is in line with recent findings that KAT2A regulates short-chain acidic acylations, including lysine succinylation and glutarylation, in histones (Bao et al., 2019; Wang et al., 2017). However, unlike EP300, KAT2A does not appear to have a deep aliphatic pocket in its active site to accommodate longer-chain acyl-CoA species. For this reason, the acyltransferase activity of KAT2A rapidly decreases with the increase of the length of the acyl group among acetyl-, propionyl-, and butyryl-CoA species (Ringel and Wolberger, 2016). It is unclear if KAT2A undergoes structural rearrangement to accommodate short-chain acidic acyl-CoA species. Hence, further structural studies will be needed to resolve the molecular mechanism.

Compartmentalized acyl-CoA metabolism is important for chromatin regulation (Trefely et al., 2020). Studies have shown that histone acylations, including acetylation, crotonylation, and succinylation, are regulated by the corresponding acyl-CoA-producing enzymes localized in the nucleus (Liu et al., 2017; Liu et al., 2021; Nagaraj et al., 2017; Sutendra et al., 2014; Wang et al., 2017; Wellen et al., 2009). In the present study, we found that, in addition to the canonical cytoplasmic localization, ACC was present in nucleus, and partly localized in nucleoli. Consequently, we observed that increasing malonylation (by orlistat treatment, overexpression of KAT2A, or knocking down of SIRT5) increased nucleolar area and rRNA expression. Recent studies have shown that nucleolar enlargement is associated with aging (Buchwalter and Hetzer, 2017; Tiku et al., 2016). In the present study, we observed that lysine malonylation and ACC expression levels were higher in older mouse brains. Hence, increased histone malonylation might be a contributing factor that leads to aging-associated nucleolar expansion. Future profiling of histone malonylation at a genome-wide scale should help provide a broader view of aging-associated epigenetic changes.

To summarize, we demonstrated that malonylation is affected by malonyl-CoA availability, and regulated by SIRT5 and KAT2A. Interestingly, we also found that inducing histone malonylation increases nucleolar size and rRNA expression. Finally, our finding of an age-related increase in malonlylation and the previously reported link between nucleolar size and aging suggest a possible role for histone malonylation in the aging process.

## Materials and Methods

### Cell culture

K562 cells were cultured in RPMI1640 medium, and HEK293T and HeLa cells were cultured in Dulbecco’s modified Eagle medium (DMEM) with 10% fetal bovine serum (FBS) and 1% penicillin–streptomycin. The cells were grown at 37°C in a humidified incubator containing 5% CO_2_. Orlistat (Selleck, Cat. #1629) and AICAR (Selleck, Cat. #S1802) were dissolved in DMSO and diluted to specific concentrations for drug treatment. For transient overexpression of KAT2A, K562 or HeLa cells were transfected with pEBB-Flag-GCN5 (Addgene, Plasmid #74784) or an empty vector using Lipofectamine 3000 (Invitrogen, Cat. #L3000), according to manufacturer’s instructions 48 h before fixation or harvest.

### Mice

Sirt5 knockout (KO) mice (B6; 129 background, https://www.jax.org/strain/012757), and wild-type C57BL/6J were purchased from the Jackson Laboratory. All animal procedures were approved by Institutional Animal Care and Use Committee at Gladstone Institutes; University of California, San Francisco; and Buck Institute for Research on Aging. Mice were housed (12-h light/dark cycle, 22°C) and given unrestricted access to water.

### Construction of CRISPRi K562 cells

K562 cells stably expressing a dCas9-KRAB fusion protein were kindly provided by Jonathan S. Weissman (Gilbert et al., 2013). pU6-sgRNA-EF1Alpha-puro-T2A-BFP (Addgene, Plasmid #60955) was used to express sgKAT and sgNC sgRNAs; and pU6-sgRNA-EF1Alpha-BSR-T2A-BFP (puromycin-resistance gene replaced by blasticidin S-resistance gene) was used to express sgSIRT5 sgRNAs. The generation of stably gene knockdown K562 cells was performed using a CRISPRi system (Gilbert et al., 2014). Briefly, double-stranded sgRNA oligos (**Table S1**) were synthesized and inserted into backbone plasmids using BstXI and BlpI restriction sites. Lentivirus expressing sgRNAs were packaged and produced using HEK293T cells with pCMV-VSV-G envelope vector (Cell Biolabs, Cat. # RV-110) and pCMV-dR8.91 packaging vector (Addgene, Plasmid #2221). K562-dCas9-KRAB cells stably expressing sgRNAs were generated upon lentiviral infection, and selected using corresponding antibiotics and fluorescence-activated cell sorting of BFP-positive cells.

### Reverse transcription and qPCR

RNA was extracted from HeLa and K562 cells using RNA STAT-60 (Fisher Scientific, Cat. #NC9489785). Reverse transcription was performed using a First-Strand cDNA Synthesis kit (New England Biolabs, Cat. #E6560) and random primers, according to manufacturer’s instructions. RNA expression levels were quantified by qPCR with a SYBR Green reagent (Thermo Fisher Scientific, Cat. #K0223). Primers for qPCR are shown in **Table S2**.

### Western blotting

Cells and tissues are lysed and homogenized with RIPA buffer (Thermo Fisher Scientific, Cat. #89900) containing protease and phosphatase inhibitor cocktail (Thermo Fisher Scientific, Cat. # PI78447), 5 M trichostatin A (Cayman Chem, Cat. #89730), and 5 mM nicotinamide (Sigma-Aldrich, Cat. #72340). Histones were extracted from cells and tissues using a total histone extraction kit (EpiGentek, Cat. #OP-0006). Before SDS-PAGE, protein samples were heated at 70℃ for 5 min. Antibodies used are listed as follows: anti-β-tubulin (Cell Signaling Technology, Cat. #2128), anti-SIRT5 (Cell Signaling Technology, Cat. #8782), anti-acetyl-lysine (Cell Signaling Technology, Cat. #9814), anti-malonyl-lysine (PTM Biolabs, Cat. #PTM-901), anti-succinyl-lysine (PTM Biolabs, Cat. # PTM-401), anti-glutaryl-lysine (PTM Biolabs, Cat. #PTM-1151), anti-ACC (Cell Signaling Technology, Cat. #3676), anti-phospho-ACC (Ser79) (Cell Signaling Technology, Cat. #3661), anti-H2B (Cell Signaling Technology, Cat. #12364), anti-KAT2A (Cell Signaling Technology, Cat. #3305), anti-HSP60 (Abcam, Cat. #ab59457), Lamin A/C (Cell Signaling Technology, Cat. #12741), COX IV (Abcam, Cat. #ab16056), and GAPDH (Cell Signaling Technology, Cat. #5174). Ponceau S solution (Sigma-Aldrich, Cat. #P7170) was used for protein staining.

### Subcellular fractionation

Subcellular fractionation was performed as described (Park et al., 2013). Briefly, HEK293T cells were gently homogenized in hypotonic buffer on ice with a dounce homogenizer till the vast majority of cells were ruptured, and the nuclei were blue stained by Trypan blue. Cells were centrifuged at 600 g for 3 min at 4°C to obtain a pellet (nuclear fraction). The supernatant was further centrifuged at 15,000 g for 10 min at 4°C to pellet mitochondria, and the supernatant was collected as the cytosolic fraction. For western blotting, equal protein amounts of each fraction were assessed.

### Immunofluorescence staining and microscopy

HeLa cells were seeded on to slide chambers (Waston Bio Lab, Cat. #192-008), and treated with DMSO or orlistat for 24 h before fixation with 4% paraformaldehyde (PFA) for 15 min at room temperature. Cells were permeabilized with PBS containing 1% BSA and 0.01% Triton X-100 for 15 min, and blocked with PBS containing 1% BSA, 5% goat serum (same source as the secondary antibodies) and 0.1% Tween 20 for 1 h at room temperature. Cells were incubated with primary antibodies at 4°C overnight. Primary antibodies used are anti-ACC (Cell Signaling Technology, Cat. #3676, 1:200 dilution) and anti-FBL (abcam, Cat. #ab4566, 1:500 dilution). Cells were incubated with secondary antibodies at room temperature for 1 h away from light. Secondary antibodies were goat anti-rabbit IgG, Alexa Fluor 555 (Invitrogen, Cat. #A-21428, 1:500 dilution) and goat anti-mouse IgG, Alexa Fluor 488 (Invitrogen, Cat. #A-11001, 1:500 dilution). Nuclei were stained with DAPI (Sigma-Aldrich, Cat. #D9542) before mounting with ProLong antifade mountant (Invitrogen, Cat. # P10144).

For confocal microscopy, Zeiss LSM 780 was used with 63x oil immersion objective at Ex. (nm) / Detection (nm) = 405/410-508 (DAPI), 488/490-570 (Alexa Fluor 488), and 561/566-697 (Alexa Fluor 555). Fluorescence intensity distribution was generated using Zeiss Zen 3.4 (blue edition). For wide-field microscopy, Keyence BZ-X800 was used with a 40x objective. 20 images were taken for each genotype and treatment group. Nucleolar percentage area was quantified using a machine-learning algorithm, where we used U-NET for image segmentation to automatically detect DAPI-stained nuclei and fibrillarin-stained nucleoli and then quantified nucleoli and nuclei area (Ronneberger et al., 2015). Several U-NET models were trained, including HeLa nuclei with four images, HeLa nucleoli with four images, HEK293T nuclei with four images, and HEK293T nucleoli with eight images. Each training session utilized data augmentation, including rotation, scaling, and brightness variation, and was performed for 1000 epochs. After training, U-NET was used to predict the nuclei and nucleoli area for each cell type. Each detected region was then scanned using a flood fill algorithm to quantify area in pixels.

### Mass spectrometry

Identification of malonylation sites in histones in mouse liver tissues was performed as reported (Nishida et al., 2015). Briefly, livers were collected from five WT and five SIRT5 KO male mice at 10 months of age. Anti-malonyl-lysine antibodies (Cell Signaling Technology, H4597 and BL13640) were used for immunoaffinity enrichment of malonylated peptides. All samples used for MS1 filtering experiments were analyzed by reverse-phase LC-ESI-MS/MS with an Eksigent Ultra Plus nano-LC 2D HPLC system connected to a quadruple time-of-flight TripleTOF 6600 mass spectrometer (MS) (SCIEX) in direct injection mode. Data sets were analyzed and searched using Mascot server version 2.3.02 (Matrix Sciences) and ProteinPilot (AB SCIEX 4.5beta) with the Paragon algorithm (4.5, 1656). All data files were searched using the SwissProt 2013_01 database with a total of 538,849 sequences but restricted to *Mus musculus* (16,580 protein sequences).

To identify malonylation sites in histones, four 18-month-old WT and four SIRT5 KO female mice were fasted for 24 h and refed for 6 h before sacrifice. Frozen brains were immersed in lysis buffer containing 8 M urea, 200 mM triethylammonium bicarbonate (TEAB), pH 8.5, 75 mM sodium chloride, 1 μM trichostatin A, 3 mM nicotinamide, and 1x protease/phosphatase inhibitor cocktail (Thermo Fisher Scientific, Waltham, MA), homogenized for 2 cycles with a Bead Beater TissueLyser II (Qiagen, Germantown, MD) at 30 Hz for 3 min each, and sonicated for 10 min. Lysates were clarified by spinning at 14,000 x g for 10 min at 4°C, and the supernatant containing the soluble proteins was collected. Protein concentrations were determined using a bicinchoninic acid protein assay (Thermo Fisher Scientific), and subsequently 5 mg of protein from each sample were brought to an equal volume using a solution of 8 M urea in 50 mM TEAB, pH 8. Proteins were reduced using 20 mM dithiothreitol in 50 mM TEAB for 30 min at 37 °C, and after cooling to room temperature, alkylated with 40 mM iodoacetamide for 30 min at room temperature in the dark. Samples were diluted four-fold with 50 mM TEAB, pH 8, and proteins were digested overnight with a solution of sequencing-grade trypsin (Promega, San Luis Obispo, CA) in 50 mM TEAB at a 1:50 (wt:wt) enzyme:protein ratio at 37°C. This reaction was quenched with 1% formic acid (FA), and the sample was clarified by centrifugation at 2,000 x g for 10 min at room temperature. Clarified peptide samples were desalted with Oasis 10-mg Sorbent Cartridges (Waters, Milford, MA). 100 μg of each peptide elution were aliquoted for analysis of protein-level changes, after which all desalted samples were vacuum dried. The 100c whole lysate were re-suspended in 0.2% FA in water at a final concentration of 1 µg/µL and stored for MS analysis. The remaining 4.9 mg of digests were re-suspended in 1.4 mL of immunoaffinity purification buffer (Cell Signaling Technology, Danvers, MA) containing 50 mM 4-morpholinepropanesulfonic acid (MOPS)/sodium hydroxide, pH 7.2, 10 mM disodium phosphate, and 50 mM sodium chloride for enrichment of posttranslational modifications. Peptides were enriched for malonylation with anti-malonyl antibody conjugated to agarose beads from the Malonyl-Lysine Motif Kit (Cell Signaling Technology, Danvers, MA). This process was performed according to the manufacturer protocol; however, each sample was incubated in half the recommended volume of washed beads. Peptides were eluted from the antibody-bead conjugates with 0.15% trifluoroacetic acid in water and were desalted using C18 stagetips made in-house. Samples were vacuum dried and re-suspended in 0.2% FA in water. Finally, indexed retention time standard peptides (iRT; Biognosys, Schlieren, Switzerland) (Escher et al., 2012) were spiked in the samples according to manufacturer’s instructions. Samples were analyzed in data-independent acquisition (DIA) mode on a Dionex UltiMate 3000 system coupled to an Orbitrap Eclipse Tribrid MS (both from Thermo Fisher Scientific, San Jose, CA). The solvent system consisted of 2% ACN, 0.1% FA in water (solvent A) and 98% ACN, 0.1% FA in water (solvent B). Peptides enriched in posttranslational modifications (4 µL) were loaded onto an Acclaim PepMap 100 C18 trap column (0.1 x 20 mm, 5 µm particle size; Thermo Fisher Scientific) over 5 min at 5 µL/min with 100% solvent A. Peptides were eluted on an Acclaim PepMap 100 C18 analytical column (75 µm x 50 cm, 3 µm particle size; Thermo Fisher Scientific) at 0.3 µL/min using the following gradient of solvent B: linear from 2–20% in 125 min, linear from 20–32% in 40 min, and up to 80% in 1 min, with a total gradient length of 210 min. For the DIA analysis, full MS spectra were collected at 120,000 resolution (AGC target: 3e6 ions, maximum injection time: 60 ms, 350–1,650 m/z), and MS2 spectra at 30,000 resolution (AGC target: 3e6 ions, maximum injection time: Auto, NCE: 27, fixed first mass 200 m/z). The isolation scheme consisted in 26 variable windows covering the 350–1,650 m/z range with a window overlap of 1 m/z (**Table S3**) (Bruderer et al., 2017).

A spectral library was generated in Spectronaut (version 14.10.201222.47784; Biognosys, Schlieren, Switzerland) from the DIA acquisitions using a *Mus musculus* UniProtKB-TrEML database (58,430 entries, 01/31/2018). BGS settings were set as defaults settings, except for the Pulsar search for which four missed cleavages were allowed and lysine malonylation was added as variable modification. The final library contained 13,304 modified peptides and 3,112 protein groups (**Table S4**). The library was further imported into Skyline-daily (version 21.1.1.223) (MacLean et al., 2010). DIA data extraction for the peptide PDPAKSAPAPK (H2B2B_MOUSE) was performed using the following parameters: up to 10 of the most intense product ions (monocharged y- and b-type ions) from ion 2 to last ion-1 per precursor ion were extracted. Resolving power was set to 30,000 at 400 m/z, and all matching scans were including. Finally, chromatographic peaks were investigated to manually adjust peak integration boundaries and remove interfered transitions (four final transitions). Signal at the peptide level was obtained by summing the corresponding transition peak areas. Statistical analysis was performed in Skyline using default settings.

## Data Availability

Raw data and complete MS data sets have been uploaded to the Center for Computational Mass Spectrometry, to the MassIVE repository at UCSD, and can be downloaded using the following link: http://massive.ucsd.edu/ProteoSAFe/status.jsp?task=a23d5ddef1b54e89b858221ce 57863ab (MassIVE ID number: MSV000089318; ProteomeXchange ID: PXD033458).

## Statistical analyses

GraphPad Prism 6 (GraphPad Software, San Diego, CA) was used for statistical analyses. Unpaired two-tailed Student’s t-test was used when two groups of data were compared. Sidak multiple comparisons test following two-way ANOVA was used when malonylation levels between WT and S5KO groups on a certain lysine site were compared, and when rRNA expression levels between WT and S5KO groups were compared. P values less than 0.05 were considered statistically significant. Optical densities of western blots were quantified using ImageJ software (Schindelin et al., 2012).

## Author contributions

RZ, BS and EV designed the experiments and analyzed the data. JB and JR performed mass spectrometry analysis. OB made sgSIRT5 HEK293T cell lines. IH performed nucleolar area quantification analysis. RZ performed the rest of the experiments. RZ and EV wrote the manuscript. All authors discussed the results and commented on the manuscript.

## Acknowledgements

We thank Dr. Xueshu Xie for discussion and suggestions. We thank Dr. Marius Walter for assistance with CRISPRi K562 cell construction; Dr. Herbert Kasler for fluorescence-activated cell sorting; Ms. Lei Wei for assistance with sample processing for mass spectrometry, Dr. Wenjuan He for providing mouse tissue samples,; Ms. Rebeccah Riley and Mr. Ryan Kwok for mouse colony maintenance; Mr. John Carroll for graphic design; Mr. Gary Howard for language editing. This study was supported by NIH grant R24DK085610 (Eric Verdin and Birgit Schilling), Glenn Foundation for Medical Research (Ran Zhang), and NCRR shared instrumentation grants 1S10 OD016281 (TripleTOF 6600, Buck Institute) and 1S10 OD028654 (Orbitrap Eclipse Tribrid, PI: Birgit Schilling).

## Supplemental Figure Legend

**Figure S1.**
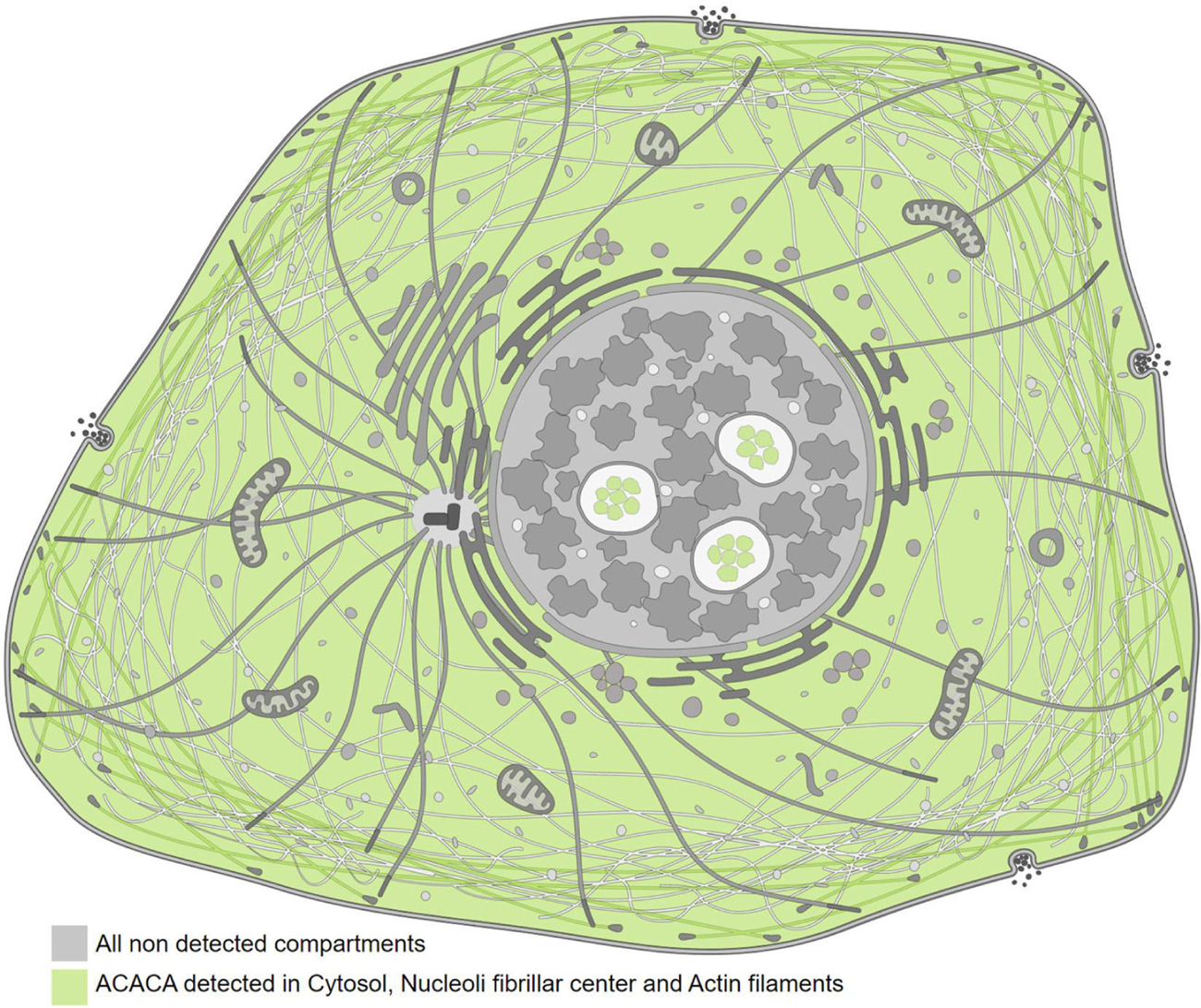
ACC1 subcellular localization (the HPA database, proteinatlas.org).

